# Angiotensin Converting Enzyme Hub Model of Peptide Flux Relating Calculated AT1 Receptor and Aldosterone Levels

**DOI:** 10.1101/2024.08.20.608830

**Authors:** Brian M. Westwood, Mark C. Chappell

## Abstract

Utilizing data from the Vitamin C, Thiamine, and Steroids in Sepsis (VICTAS) Trial, this hub model was developed to limit seventeen Renin-Angiotensin-Aldosterone System (RAAS) components as three entrance and four exits, to facilitate the calculation of a model as one egress unknown, the angiotensin type 1 (AT1) receptor. Following previous evidence relating renin levels to mortality, this study found controls were more like sepsis patients with levels < renin quartile 1 (Q1) for calculated AT1, while more like sepsis patients with renin levels > quartile 3 (Q4) for measured aldosterone levels. Additionally differential discrete correlate summation (DCS) analysis indicates AT1, aldosterone and renin as major hub nodes in this independent DCS model of metabolic data inputs.

## INTRODUCTION

Under classical physiologic conditions, angiotensin converting enzyme (ACE, EC 3.4.15.1) is a critical hub node for the RAAS and the conservation of the precursor angiotensinogen (Ao) consumption for a direct chain of conversions to angiotensin-(1-5) [Ang-(1-5)]. Angiotensin I [Ang-(1-10) or Ang I] is generated by renin [EC 3.4.23.15] from Ao, then ACE yields the first N-terminally intact bioactive peptide (BiP) in this cascade, angiotensin II [Ang-(1-8) or Ang II], which is converted to the next BiP, angiotensin-(1-7) [Ang-(1-7)], then ACE generates Ang-(1-5) (Chappell, 2016) (Figure 1). Both bioactive peptides are degraded by dipeptidyl peptidase 3 (DPP3, EC:3.4.14.4).

**Figure 1.**
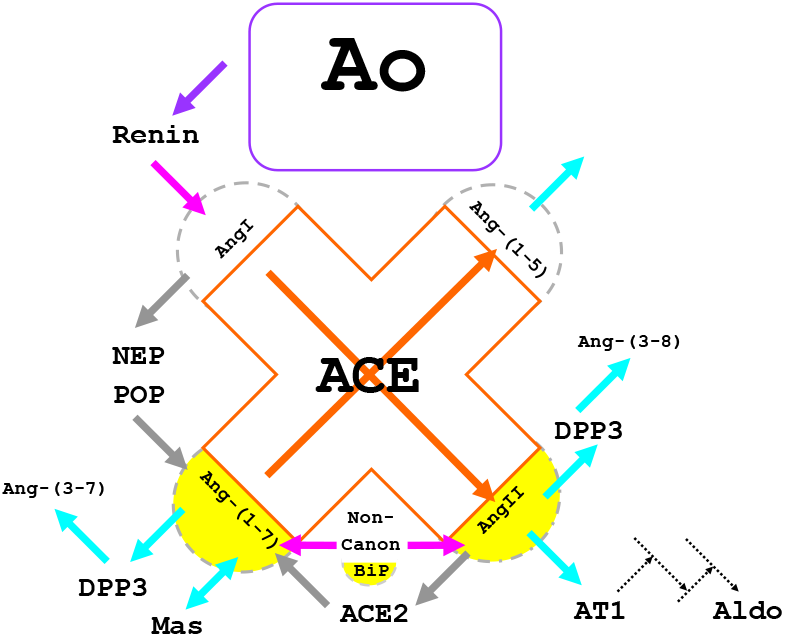
Simple model for ACE hub with the control volume extended to substrates and products, for development of conservation equation mathematical model for RAAS components.s

We recently completed a post-hoc analysis of the RAAS in a subset of patients from the VICTAS septic shock trial that revealed a strong association of renin to mortality (Busse et al., 2023, Chappell et al., 2024, Chappell et al., 2025). As plasma samples were not collected with a renin inhibitor, both intact and total Ao levels were measured for comparison, acknowledging intact levels may be lower than in-vivo (intact Ao was used in the conservation model calculations). To assess this system, we developed a conservation model that incorporated the internalization of the Ang II type 1 (AT1) receptor (Zhuo et al., 2002) and utilized the RAAS data from the VICTAS trial, which may be applied to other situations to calculate an unknown member of the RAAS (Westwood et al., 2008).

## METHODS

If we consider ACE and its substrates and products from Figure 1 as an extended hub control volume (ACE hub anastomoses, gray arrows), we can write the conservation equations [this model assumes that under classical physiologic conditions Ang I generated by renin is converted to Ang II by ACE, then directly converted to Ang-(1-7) by ACE2 [EC 3.4.17.23] and is acted upon by ACE to generate Ang-(1-5)]. There are three additional egression points in the ACE hub linear model for BiP via DPP3, as well as AT1 receptor uptake of Ang II, and non-canonical BiP entrance. Under the pseudo-steady state hypothesis (PSSH), we can set the accumulation term to zero.

Given the ACE hub (Figure 1), the conservation of Ang I, Ang II and Ang-(1-7) can be described:

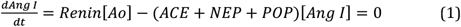

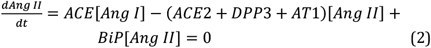

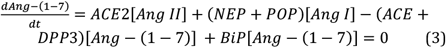

ACE hub eqn. 4 = eqns. 1 + 2 + 3

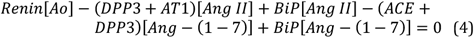

Enzyme concentrations were used as proxy for rate constants. The PSSH yields the key to calculating AT1 levels:

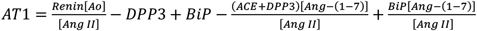

DPP3 has a substantially lower efficacy of kcat/Km (s^-1^μM^-1^) of 0.03 and 0.07 for Ang II and Ang-(1-7) (Jha et al.,2020), respectively than renin has for Ao of 0.496 (Cumin et al., 1987) relative to 2.2 of ACE for Ang-(1-7) (Chappell et al., 1998). However, DPP3 is a major sink in the system, providing two egress points; as such DPP3 extreme outliers (above 3xIQR+Q3, upper outer fence, UOF) which range to >10 fold higher than UOF (Tukey, 1977) were Winsorized to the UOF value (Tukey, 1962), for AT1 calculations. Bioactive peptides from noncanonical production were set to a constant of 45nM (concentration, same as enzyme rate and receptor affinity proxies) to achieve all positive AT1 calculated values. These BiP source activities are modulated for each subject individually on both Ang II and Ang-(1-7) concentrations.

Quartile metrics based on subjects in all five groups.

The model input variables were analyzed by differential discrete correlate summation (DCS). Briefly |log correlation ratios| were tabulated for low renin (<Q2) relative to high renin (>Q2) correlations for all variable combinations (Westwood et al., 2006 and Bronson et al., 2022).

All calculations were performed in Excel (Microsoft Corp., WA), Prism (GraphPad Software, MA) was used for plotting data in figures and statistical analyses.

## RESULTS

IQR-normalization vs. ACE hub model variables (Figure 2). IQR-score calculated for each group individually. If |IQR-score| > half of group total, highlighted in purple for below IQR and orange for above IQR (Table 1).

**Table 1.** IQR-scores. Orange highlight if at least half more than IQR and purple if at least half less than IQR.

| gp | n | Ren pM | ACE nM | ACE2 nM | DPP3 nM | All pM | A7 pM | TotAo nM | Ao-nM | ALDO pM | AT1 |
| --- | --- | --- | --- | --- | --- | --- | --- | --- | --- | --- | --- |
| Controls | 20 | -13 | 12 | -8 | -18 | 6 | -13 | 4 | 6 | 2 | -13 |
| Q1 | 14 | -10 | -3 | 1 | 5 | -3 | 1 | 7 | 9 | -10 | -3 |
| Q2 | 21 | 0 | -4 | 1 | 4 | -5 | 0 | -1 | 0 | 1 | 2 |
| Q3 | 19 | 7 | -2 | 1 | 2 | 1 | 4 | 1 | -3 | 3 | 7 |
| Q4 | 16 | 16 | -3 | 5 | 7 | 1 | 8 | -11 | -12 | 4 | 7 |

**Figure 2.**
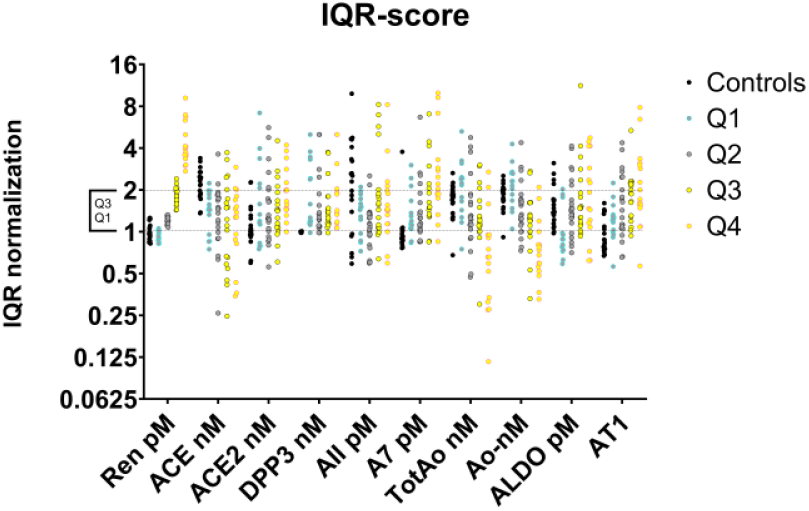
Group independent IQR-normalization vs. ACE hub model variables. Controls represented by black, Q1 cyan/gray (border/fill), Q2 black/gray, Q3 black/yellow and Q4 by magenta/yellow circles.

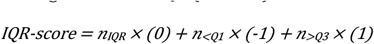

Fisher’s exact test (p = 0.0014) and area under the curve [AUC = 0.8089 (0.6921,0.9257 - 95% confidence interval), p < 0.0001] for receiver operating characteristic (ROC) curves (Fawcett, 2006) were calculated for |Q4-Q1| for less than controls vs. at least controls measured aldosterone values relative to increasing calculated AT1 values (Figure 3).

**Figure 3.**
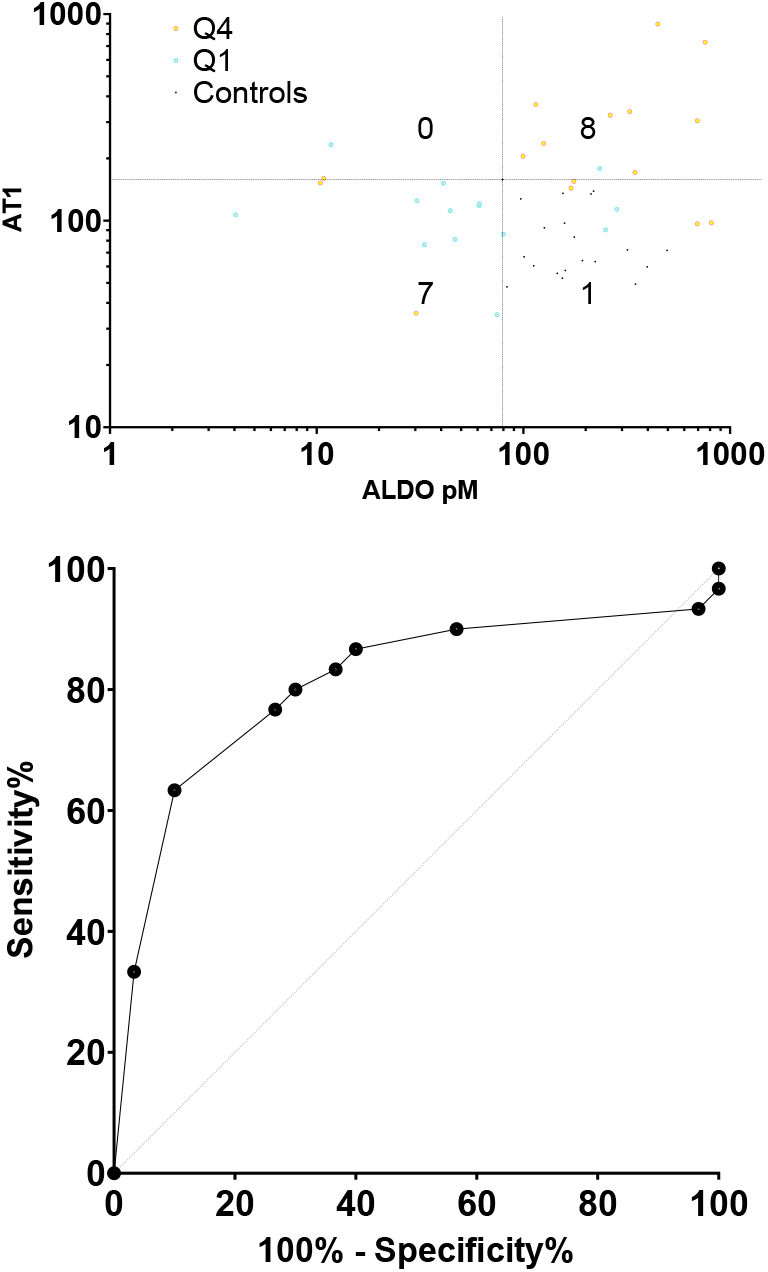
Top panel, values for |Q4-Q1| displayed in each quadrant for AT1 vs. Aldo. Controls are black dots, contained entirely in fourth quadrant. Q4 (magenta) and Q1 (cyan) circles mostly in first and third quadrants, respectively. Bottom panel, ROC curve for |Q4-Q1| for less than controls vs. at least controls measured aldosterone values relative to increasing calculated AT1 values.

## DISCUSSION

Autonomic stress is complicated (Goldstein, 2021). Modeling the RAAS may prove advantageous, given that Ang II has been found to be beneficial in experimental models of sepsis and in clinical care (Leisman et al., 2022, Garcia et al., 2024, Flannery et al., 2022). Given the benefits of RAAS inhibitions for chronic diseases, and the possibility of small interfering ribonucleic acid treatments that can last for weeks to months (Cruz-López et al., 2022), perspective is important. AT1 as a pivot with aldosterone for the state of these septic shock patients around a control cohort, evokes the utility of modeling this system. Additionally, DCS indicates AT1 and aldosterone along with renin as hub nodes in a naïve network of this data. Figure 4 shows the overlay of differential DCS edges with > 10-fold change in p-value between low and high renin ([Q1,Q2]:[Q3,Q4]) sepsis patients (from figure 5). This model may represent a seed method for the description of the complex interplay between not only this select catalog of RAAS components, but also may be generalized as a paradigm to explore greater utility in more complex systems.

**Figure 4.**
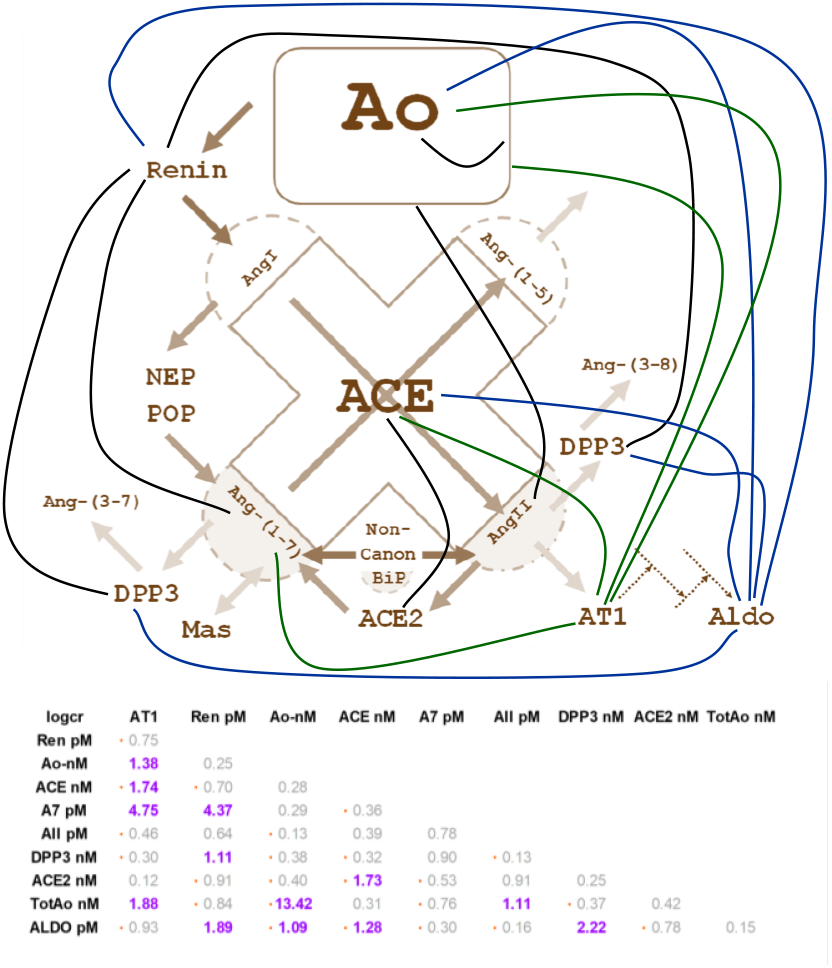
ACE hub pathway model with overlay of differential DCS system diagram. AT1 edges are green, and aldosterone edges are blue. Table inlay lists maximum log correlation ratios (logcr) for either linear or log-log (orange dot) correlations. All edges in diagram are logcr > 1 (more than 10x difference in correlation p-values between low and high renin) are purple in table.

**Figure 5.**
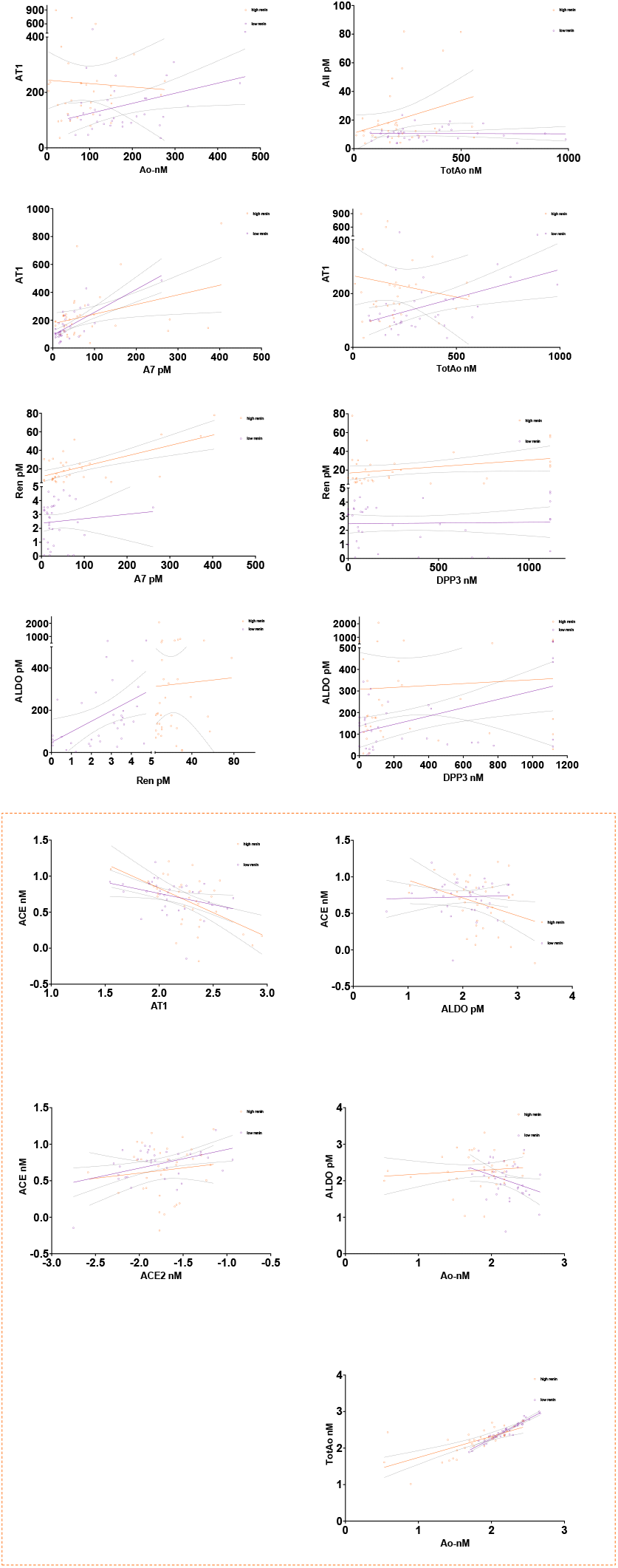
DCS plots. Linear and log-log (orange border, values log_10_[*]) regressions with 95% confidence intervals for low (purple) and high (orange) renin ([Q1,Q2]:[Q3,Q4]) sepsis patients for logcr > 1 correlations in figure 4.

## ACKNOWLEDGMENT

The authors acknowledge support from the National Heart, Lung, and Blood Institute (R01HL146818) and the Cardiovascular Sciences Center (pilot award).

## ABBREVIATIONS

VICTAS: Vitamin C, Thiamine, and Steroids in Sepsis
ACE: angiotensin converting enzyme
RAAS: Renin-Angiotensin-Aldosterone System
AT1: angiotensin type 1 receptor
DCS: differential discrete correlate summation analysis
Q*: quartile *
Ao: angiotensinogen
Ang-(1-5): angiotensin-(1-5)
Ang I or Ang-(1-10): Angiotensin I
Ang II or Ang-(1-8): angiotensin II
Ang-(1-7): angiotensin-(1-7)
ACE hub: ACE and its substrates and products as a hub control volume extended
BiP: bioactive peptides
PSSH: pseudo-steady state hypothesis
W: Winsorized
AUC: area under the curve
ROC: receiver operating characteristic.

## REFERENCES

Chappell MC. Biochemical evaluation of the renin-angiotensin system: the good, bad, and absolute? Am J Physiol Heart Circ Physiol. 2016 Jan 15;310(2):H137–52. doi: 10.1152/ajpheart.00618.2015. Epub 2015 Oct 16. PMID: 26475588; PMCID: PMC4796631.

Busse, Laurence W.; Schaich Christopher L.; Chappell Mark C.; McCurdy Michael T.; Staples Erin M.; Ten Lohuis, Caitlin C.; Hinson Jeremiah S.; Sevransky Jonathan E., MHS1,2; Rothman, Richard E.; Wright David W.; Martin Greg S.; Khanna Ashish K. Association of Active Renin Content With Mortality in Critically Ill Patients: A Post hoc Analysis of the Vitamin C, Thiamine, and Steroids in Sepsis (VICTAS) Trial*. Critical Care Medicine 52(3):p 441–451, March 2024. | DOI: 10.1097/CCM.0000000000006095

Chappell MC, Schaich CL, Busse LW, Martin GS, Sevransky JE, Hinson JK, Khanna AK; Vitamin C, Thiamine, Steroids in Sepsis (VICTAS) Investigators. Stronger association of intact angiotensinogen with mortality than lactate or renin in critical illness: posthoc analysis from the VICTAS trial. Crit Care. 2024 Oct 14;28(1):333. doi: 10.1186/s13054-024-05120-w. PMID: 39402593; PMCID: PMC11472595.

Chappell MC, Schaich CL. Busse LW, Files D. Clark, Martin GS, Sevran-sky JE, Hinson JK, Rothman R, Khanna AK; Vitamin C, Thiamine, Ster-oids in Sepsis (VICTAS) Investigators. Higher Circulating ACE2 and DPP3 but Reduced ACE and Angiotensinogen in Hyperreninemic Sepsis Patients Clinical Sciences (London), in press, 2025.

Zhuo J. L., Imig J. D., Hammond T. G., Orengo S., Benes E. and Navar L. G. Ang II Accumulation in Rat Renal Endosomes During Ang II-Induced Hypertension. Hypertension 2002 Vol. 39 Issue 1 Pages 116–121. DOI: 10.1161/hy0102.100780

Westwood BM, Hossam S, Chappell MC. Modeling of Angiotensin Peptide Metabolism in Renal Proximal Tubules. ASME 2008 Summer Bioengineering Conference. 2008 June; SBC2008-190990:181. doi:<https://asmedigitalcollection.asme.org/SBC/proceedings-abstract/SBC2008/43215/181/287274>.

Tukey, John W.: Exploratory Data Analysis. Addison-Wesley Publishing Company Reading, Mass. — Menlo Park, Cal., London, Amsterdam, Don Mills, Ontario, Sydney 1977, XVI, 688 S.

Tukey, John W. “The Future of Data Analysis.” The Annals of Mathematical Statistics; 33(1) 1–67, March, 1962. 10.1214/aoms/1177704711

Jha S, Taschler U, Domenig O, Poglitsch M, Bourgeois B, Pollheimer M, Pusch LM, Malovan G, Frank S, Madl T, Gruber K, Zimmermann R, Macheroux P. Dipeptidyl peptidase 3 modulates the renin-angiotensin system in mice. J Biol Chem. 2020 Oct 2;295(40):13711–13723. doi: 10.1074/jbc.RA120.014183. Epub 2020 Jun 16. PMID: 32546481; PMCID: PMC7535908.

Cumin F, Le-Nguyen D, Castro B, Menard J, Corvol P. Comparative enzymatic studies of human renin acting on pure natural or synthetic substrates. Biochim Biophys Acta. 1987 May 27;913(1):10–9. doi: 10.1016/0167-4838(87)90226-3. PMID: 3555621.

Chappell MC, Pirro NT, Sykes A, Ferrario CM. Metabolism of angiotensin-(1-7) by angiotensin-converting enzyme. Hypertension. 1998 Jan;31(1 Pt 2):362–7. doi: 10.1161/01.hyp.31.1.362. PMID: 9453329.

Fawcett, T., An introduction to ROC analysis. Pattern Recognition Letters, 2006. 27(8): p. 861–874.

Westwood B and Chappell M. 2006. Application of correlate summation to data clustering in the estrogen- and salt-sensitive female mRen2.Lewis rat. In Proceedings of the 1st international workshop on Text mining in bioinformatics (TMBIO ‘06). Association for Computing Machinery, New York, NY, USA, 21–26. 10.1145/1183535.1183542

Bronson S, Westwood B, Cook K, Emenaker N, Chappell M, Roberts D, SotoPantoja D. Discrete Correlation Summation Clustering Reveals Differential Regulation of Liver Metabolism by Thrombos-pondin-1 in Low-Fat and HighFat Diet-Fed Mice. Metabolites 2022, 12, 1036. 10.3390/metabo12111036

Goldstein DS. Stress and the “extended” autonomic system. Auton Neurosci. 2021 Dec;236:102889. doi: 10.1016/j.autneu.2021.102889. Epub 2021 Oct 2. PMID: 34656967; PMCID: PMC10699409.

Leisman D.E., Privratsky J.R., Lehman J.R., Abraham M.N., Yaipan O.Y., Brewer M.R., Nedeljkovic-Kurepa A., Capone C.C., Fernandes T.D., Griffiths R., Stein W.J., Goldberg M.B., Crowley S.D., Bellomo R., Deutschman C.S., Taylor M.D., Angiotensin II enhances bacterial clearance via myeloid signaling in a murine sepsis model, Proc. Natl. Acad. Sci. U.S.A. 10.1073/pnas.2211370119 (2022).

Garcia, B., ter Schiphorst, B., Santos, K. et al. Inhibition of circulating dipeptidyl-peptidase 3 by procizumab in experimental septic shock reduces catecholamine exposure and myocardial injury. ICMx 12, 53 (2024). 10.1186/s40635-024-00638-3

Flannery AH, Kiser AS, Behal ML, Li X, Neyra JA. RAS inhibition and sepsis-associated acute kidney injury. J Crit Care. 2022 Jun;69:153986. doi: 10.1016/j.jcrc.2022.153986. Epub 2022 Jan 24. PMID: 35085853; PMCID: PMC9727727.

Cruz-López EO, Ye D, Wu C, Lu HS, Uijl E, Mirabito Colafella KM, Danser AHJ. Angiotensinogen Suppression: A New Tool to Treat Cardiovascular and Renal Disease. Hypertension. 2022 Oct;79(10):2115–2126. doi: 10.1161/HYPERTENSIONAHA.122.18731. Epub 2022 Jul 29. PMID: 35904033; PMCID: PMC9444253.

